# A role for the *Stentor* syntaxin protein in post-wound cell survival

**DOI:** 10.1101/2025.11.04.686591

**Authors:** Ambika V. Nadkarni, Ulises Diaz, Kevin S. Zhang, Sindy KY Tang, Wallace F Marshall

## Abstract

Wound healing is an essential biological process that occurs both in tissues and single cells. In free-living single-celled ciliates such as *Stentor coeruleus*, rapid wound healing is necessary to repair breaches to the plasma membrane, where any delays represent the difference between life and death. In order to discover novel molecular pathways that are important for healing in *Stentor*, we carried out a targeted RNA interference-based perturbation genetic screen combined with microsurgical wounding using a microfluidic guillotine to introduce reproducible bisection wounds. We identified a *Stentor* syntaxin gene that was necessary for cell survival, particularly post-wounding, with only ∼37% of the syntaxin-deficient cells surviving compared with ∼98% of the control cells. Syntaxin-deficient cells were more susceptible to hyposmotic shock and became increasingly vacuolated in the hours post-wounding, eventually leading to cell death. Osmotic stabilization of the cells during and after bisection partially restored the post-wound survival in knockdown cells. These results support the interpretation that syntaxin-deficient cells lack essential membrane fusion machinery, which manifests in vacuolar defects, and are deficient in maintaining osmotic homeostasis necessary for their survival post-wounding. This study provides a template for the discovery of new wound healing biology in emerging model systems.

**Significance:**

- *Stentor* is a single-celled ciliate with prodigious healing ability, yet the molecular mechanisms by which the cell rapidly heals wounds are not fully characterized.
- Through a targeted microfluidics genetic screen, we identified a novel syntaxin-like protein that is essential for *Stentor* survival post-wounding.
- Syntaxin-deficient cells were more susceptible to osmotic shock. Wounding cells in osmotically balanced conditions reduced the degree of hyposmotic stress associated with wounding and enabled more cells to survive. Our discovery highlights the rebalancing of intracellular osmolarity as an important step in wound repair in freshwater-dwelling organisms.

## Introduction

Wound healing is a universal biological response commonly examined at the tissue level. However, wound healing also occurs in single cells, particularly those that sustain damage during normal function, such as muscle cells and neurons (Abreu-Blanco *et al*., 2011; Tang and Marshall, 2017). A rapid response to cellular wounding is especially important in free-living, single-celled ciliate organisms such as *Stentor coeruleus,* where any delays represent the difference between life and death. *S. coeruleus* is well known for its ability to heal and regenerate from drastic wounds, including from a fragment as small as 1/27th of the original cell volume (Lillie, 1896). *Stentor coeruleus* also heals bisection wounds at ∼ 8–80 μm^2^/s, representing one of the fastest healing rates in investigated biological systems (Zhang *et al*., 2021). This extreme ability to heal wounds in *Stentor* thus provides a powerful avenue for the discovery of new healing mechanisms.

In eukaryotic systems, wounding repair is initiated by the influx of calcium ions, which trigger molecular pathways that promote membrane resealing. Membrane dynamics occur in conjunction with the remodeling of the underlying cytoskeletal network (Abreu-Blanco *et al*., 2011; Sonnemann and Bement, 2011). In systems such as *Xenopus* oocytes and *Drosophila* syncytial embryos, a circumferential actomyosin ring enables wound closure through a purse-string mechanism (Bement *et al*., 1999; Abreu-Blanco *et al*., 2011). SNARE-dependent exocytic machinery is also necessary for wound healing, as demonstrated in systems such as starfish oocytes, sea urchin eggs and fibroblasts (Bi *et al*., 1995; Miyake and McNeil, 1995). Application of botulinum neurotoxins to fertilized sea urchin eggs and 3T3 fibroblasts inhibited vesicle fusion and interfered with wound healing (Steinhardt *et al*., 1994; Bi *et al*., 1995). However, it is not known whether these mechanisms operate to close wounds in *Stentor*.

The study of wound healing and regeneration in *Stentor* has historically been carried out via microsurgery and grafting techniques (Tartar, 1961). Direct imaging of membrane and cytoskeletal components in *Stentor* remains difficult because methods for transgene expression have not yet been implemented, and *Stentor’s* robust efflux system sequesters small-molecule dyes in intracellular vesicles.

To address these challenges, here we carried out an RNA-interference based genetic screen combined with a microfluidic guillotine capable of introducing reproducible bisection wounds in a high throughput manner to identify molecular pathways important for wound healing in *Stentor*. Using this approach, we identified a *Stentor* syntaxin that was necessary for cell survival post-wounding.

Syntaxins are components of the SNARE complex (Rothman, 1994) that enable vesicle fusion (Bennett *et al*., 1992). Syntaxins, a family of transmembrane proteins, contain a characteristic ∼60-residue, coiled-coiled domain known as the SNARE domain (Teng *et al*., 2001). Syntaxins are present in all eukaryotes and mediate vesicle fusion in a variety of cellular compartments such as the plasma membrane, ER, Golgi and endosomes (Teng *et al*., 2001). Different syntaxins localize to various cellular membranes and confer specificity of vesicle fusion. Syntaxins are therefore critical for a variety of cellular processes such as neurotransmitter release, insulin release and receptor trafficking (Bennett *et al*., 1992; Manickam *et al*., 2011; Chen *et al*., 2014; Kang *et al*., 2022). Due to their importance in exocytosis, syntaxins have also been implicated in wound healing in systems such as mouse skeletal muscle cells (syntaxin 4), the *C. elegans* epidermis (syntaxin 2) and squid giant axons (Detrait *et al*., 2000; Meng *et al*., 2020; Chen and Michele, 2025). Syntaxin proteins have been studied in other ciliates within the context of secretion and exocytosis. They have been shown to be important in the biogenesis of secretory organelles in *Tetrahymena* and the exocytosis of Ɣ-amino butyric acid in *Paramecium* (Dacks and Doolittle, 2004; Kissmehl *et al*., 2007; Ramoino *et al*., 2010; Kaur *et al*., 2017).

A role for syntaxin proteins in *Stentor* has not yet been described. We therefore carried out a domain and phylogenetic analysis of the syntaxin protein. We further identified the role of syntaxin in post-wound survival and adaptation to osmotic challenges in *Stentor*. Our discovery highlights an important adaptation of freshwater ciliates to their environments.

## Results and discussion

### An RNA interference-based survival screen for wound healing

The free-living ciliate *Stentor coeruleus* is successful at healing extreme wounds, but the molecular pathways by which this occurs are not known (Slabodnick *et al*., 2013). We devised an RNA interference-based survival screen for wound-healing (Fig. 1A). Candidate genes were selected from studies of wound healing in other biological systems, or from previous transcriptomics studies of *Stentor* (Supplementary Table 1). Gene pathways were perturbed using RNA interference (RNAi), and cells were wounded by bisection through a microfluidic guillotine as previously described (Slabodnick *et al*., 2014; Blauch *et al*., 2017; Slabodnick, 2019; Koppaka *et al*., 2021; Zhang *et al*., 2021). Cell survival was visually determined by observing cell and ciliary movement at t = 0 and 12 hours post-wounding. *S. coeruleus* was insensitive to the perturbation of several candidate genes that have been shown to be important in other systems. Survival was unaffected by perturbation of homologs of actin (Stecoe_7446) and Cdc42 (Stecoe_21574), implicated in actin-based purse-string closure in many systems including *Xenopus* oocytes (Supplementary Fig. S1) (Benink and Bement, 2005).

**Figure 1.**
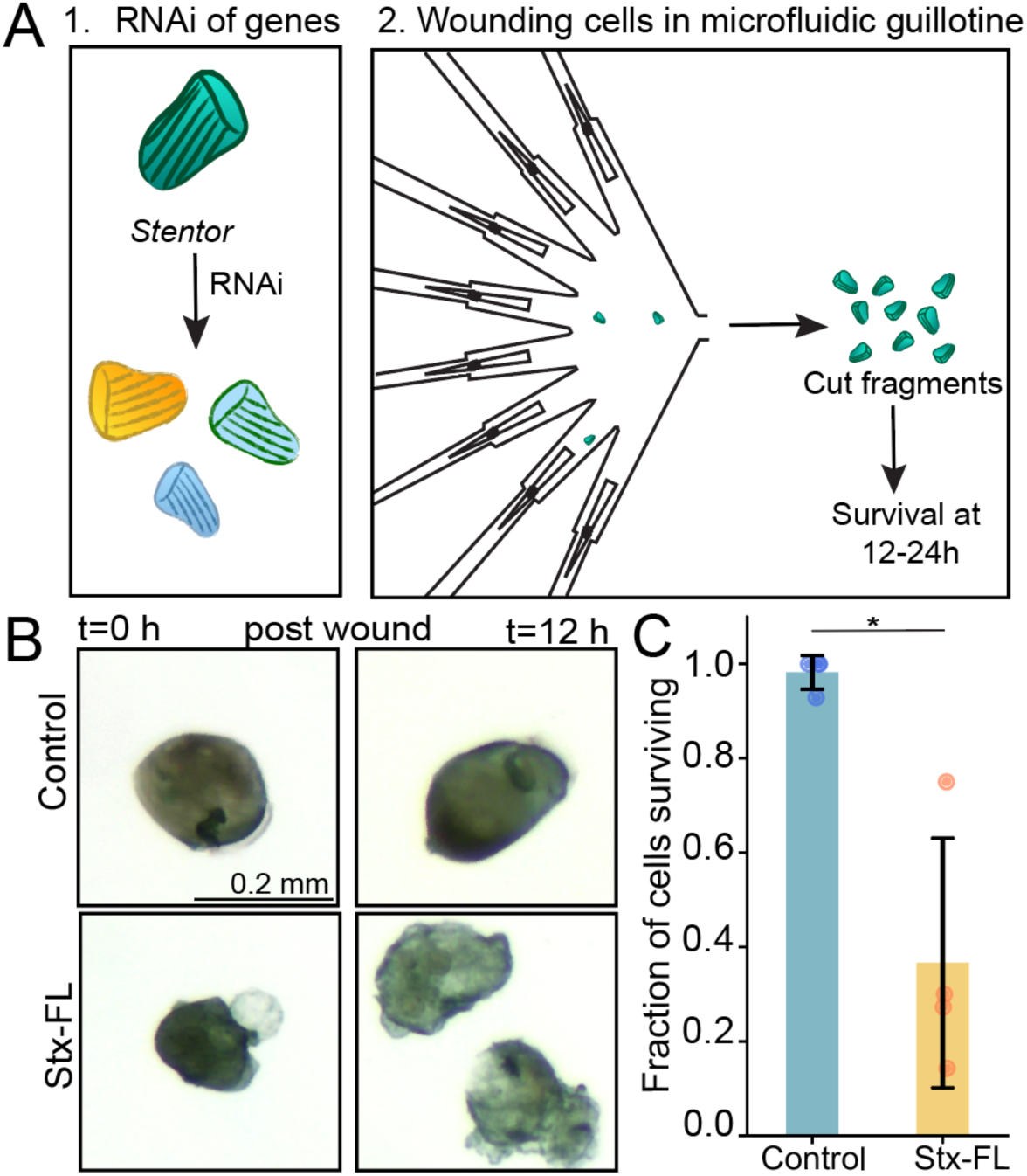
Microfluidic RNAi screen identified syntaxin as necessary for post wound cell-survival. (A) Schematic of RNAi screen.1. Left panel - Candidate genes in *Stentor coeruleus* were knocked down by feeding-based RNA interference. 2. Right panel - survival of knocked down cells was assayed by high-throughput bisection through a microfluidic guillotine at 12-24 hours post-bisection. (B) Top panels show Control cells knocked down for the gene Lf4 that did not impact cell survival post bisection at 0 h (left panels) or 12 h (right panels) after wounding. Bottom panels show the identical time point for cells knocked down for gene Stecoe_25434 (Stx-FL RNAi). (C) Quantitation of cell survival at 12 h after wounding showed the fraction of cells surviving at 12 hours was 0.982 and 0.366 for control and RNAi cells respectively (S.D. 0.036 and 0.265, p = 0.0265 by the non-parametric, two-sided Mann Whitney U-test). Points on the graph indicate the survival metrics for individual experiments. n=4 independent experiments, avg. 10 cell fragments per experiment.

Cells were also insensitive to perturbations of candidate homologs involved in membrane fusion and vesicle patching in other systems such as the C2 domain Ferlin protein (Stecoe_31310) and annexin (Stecoe_18601) (Bansal *et al*., 2003; McNeil *et al*., 2006). We also tested the role of “bisection-specific” genes in our survival assay. These genes were upregulated when cells were bisected, but not when the cells were subjected to non-wounding procedures such as sucrose-shock, which typically causes selective shedding of the mouth or oral apparatus (Sood *et al*., 2022). Among these genes, knockdown of Stecoe_168 encoding major facilitator protein 1 (MFS1) caused post-wounding survival defects. Major facilitator proteins are transmembrane transporters that mediate the transport of a variety of substrates (Supplementary Fig. S1) (Pao *et al*., 1998) but inferring substrate specificity of MFS1 from the protein sequence is beyond the scope of the current study.

We observed the most consistent and statistically significant survival defect when we inhibited a syntaxin-like protein, gene Stecoe_24534 (hereafter referred to as Stx) (Fig. 1B). Gene Lf4 was employed as a control that did not impact cell survival, as utilized previously (Slabodnick *et al*., 2014). Twelve hours post-wounding (i.e., a bisection cut), only about 37% of syntaxin (full-length)-deficient (Stx-FL RNAi) cells survived, compared with about 98% of control cells surviving (p = 0.0265, Mann Whitney U-test; n = 4, where n represents the number of independent experiments, see Methods) (Fig. 1C). Despite the limited replicates, the effect was statistically significant. Stx-FL RNAi cells appeared to lose membrane integrity with signs of rupture and disintegration (Fig. 1B, bottom right panel).

### Investigating the domain structure and phylogeny of *Stentor* syntaxin

Domain analysis of the Stecoe_24534 gene sequence revealed that the protein contained the syntaxin domain (PF 00804), a t-(or target)-SNARE domain (SM 00397) and a SNARE domain (PF 05739) (Fig. 2A). The protein was 98.61% identical (283 out of 287 amino acid residues) to the homolog protein Stecoe_1463 by a BLAST search. The two genes shared 93% sequence identity with 5-10 stretches of contiguous 20-50 base pairs without a single mismatch. Therefore it is reasonable to assume that our RNAi construct targeted both proteins. All uncovered features of the *S. coeruleus* Stx protein will also likely apply to this sister protein 1463.

**Figure 2.**
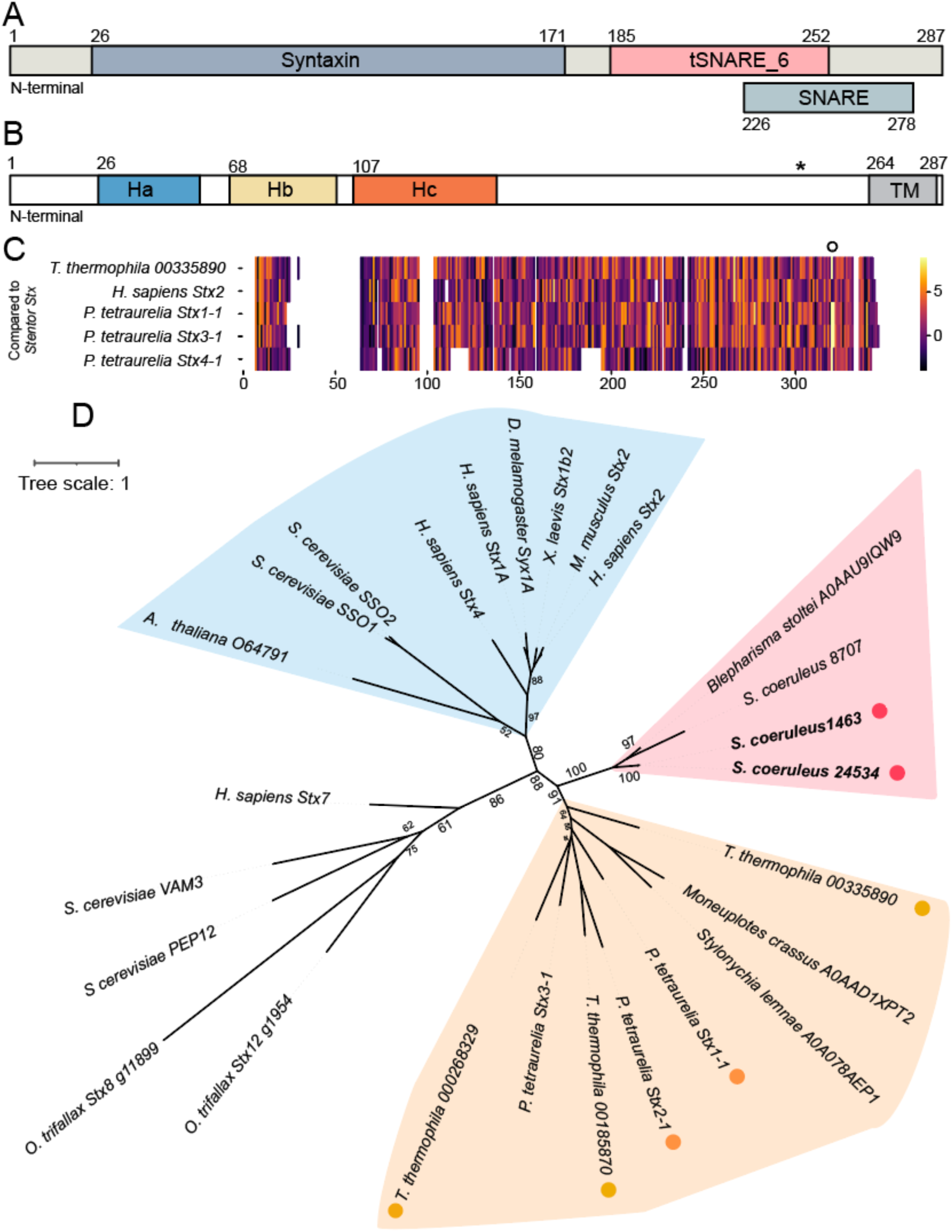
Domain structure and phylogeny of *Stentor* syntaxin. (A) InterPro domain analysis of the Stx protein revealed the presence of the syntaxin domain (slate blue), the tSNARE_6 domain (pink) and a conserved SNARE domain (light blue). Numbers indicate the amino acid residues of the protein. (B) Estimated positions of the regulatory helices Habc and the transmembrane (TM) domain in the *Stentor* Stx protein by multiple sequence alignment (msa). The conserved Q residue is also indicated (*). (C) Heat map of the BLOSUM62 similarity score of the msa of closely homologous syntaxin proteins from organisms (indicated), showing the conserved cysteine residues (yellow bars, hollow dot) at the C-terminus. (D) An unrooted phylogenetic tree of syntaxin proteins from organisms as indicated. Pink dots indicate *Stentor* Stx proteins. Homologs in *Tetrahymena thermophila (*yellow*) and Paramecium tetraurelia (*orange dots) are indicated. The pink region shows clustering of syntaxins from the subphylum Postciliodesmatophora, class Heterotrichae. The yellow region encompasses ciliates from the subphylum Intramacronucleata. Blue region shows well conserved relationships between exocytic human, fly, mouse, frog and yeast syntaxins. Numbers at the nodes indicate confidence of observations (bootstrap values). Tree scale is a measure of evolutionary change per unit length.

Syntaxins are known to contain three helices (Ha, Hb, Hc) at their N-terminal domains that are regulatory in nature (Fernandez *et al*., 1998; Lerman *et al*., 2000). We utilized previous structural work and multiple sequence alignment of syntaxin proteins to estimate the position of the 3 helices in the *Stentor* Stx protein (Fig. 2B, Supplementary Fig. S2A) (Fasshauer *et al*., 1998; Fernandez *et al*., 1998). *Stentor* syntaxin also contains a highly conserved Q (glutamine) residue present in the ionic inner core of the synaptic fusion complex. Mutation of this residue impairs fusion complex function (Fasshauer *et al*., 1998; Scales *et al*., 2001). A multiple sequence alignment of closely related syntaxin proteins from *Stentor coeruleus, Tetrahymena thermophila* and *Paramecium tetraurelia* showed the presence of this residue in all proteins (Supplementary Fig. S2B, S2C).

We constructed a heat-map of amino acid residue similarity across close homologs of *Stentor* Stx (identified in Fig. S2) using the BLOSUM similarity matrix (Fig. 2C, Supplementary Fig. S2B). A conserved pair of cysteine residues was present at the C-terminal of the Stx proteins (Henikoff and Henikoff, 1992). *Paramecium tetraurelia* Stx1-1 and Stx3-1, which have been shown to localize at the plasma membrane, also contained this dual cysteine motif, whereas *P. tetraurelia* Stx4, thought to function in transcytosis, contained one cysteine at this position (Kissmehl *et al*., 2007). *Tetrahymena thermophila* syntaxin 00335890 also contained one cysteine at this position, with the other residue being methionine. This pair of cysteines is also present in the neuronal human Stx1A protein transmembrane domain (C271, C272) and functions as a putative palmitoylation site that could anchor the protein into the membrane or lipid rafts (Masaki *et al*., 1998; Suga *et al*., 2003; Kang *et al*., 2008; Vardar *et al*., 2022; An *et al*., 2025). Due to the presence of a chain of aspartate residues present in the human Stx1a which is absent in the *Stentor* Stx, the cysteines did not align with one another in a multiple sequence alignment (Supplementary Fig. S2A). Mutation of these cysteine residues in Stx1a has been found to affect the kinetics of neurotransmitter release (Vardar *et al*., 2022; An *et al*., 2025). These residues could affect the conformation of the fusion complex or the kinetics of vesicle fusion (Vardar *et al*., 2022; An *et al*., 2025).

Next, we generated a phylogenetic tree to gain insight into the evolutionary relationship between *Stentor* Stx protein and syntaxin proteins of interest from other organisms. Because we only considered a limited number of syntaxin proteins (i.e., 26), our tree was “unrooted” and did not make conclusions about an evolutionary timeline or direction of descent, but only emphasized evolutionary relationships based on sequence similarity (Fig. 2D). *Stentor* Stx protein (gene 24534) was most related to its sister protein (gene 1463).

*Stentor* falls within the class Heterotrichea of ciliates and we found that *Stentor* Stx clusters with other heterotrich syntaxins (Fig. 2D, pink region) (Lynn, 2008; Gao *et al*., 2016). Within the other major subphylum of ciliates (Intramacronucleata) we saw extensive inter-relatedness of syntaxins (Fig. 2D, yellow region), with syntaxins in the class Spirotrichea (genus *Stylonychia*, *Moneuplotes*) apparently emerging from a common ancestor. Clustering of proteins within a subphylum increased our confidence in the validity of our phylogenetic analysis. Relationships between other eukaryotes (human, mouse, fly, yeast) obeyed current phylogenetic models (Burki, 2014). Notably, syntaxins with defined roles in exocytosis or localization at the plasma membrane appeared related (Teng *et al*., 2001), such as the yeast proteins SSO1&2 and human mouse and fly syntaxins 1,2 and human syntaxin 4 (Fig. 2D, blue region). Putative endosomal/vacuolar syntaxins were found to cluster separately from the plasma membrane syntaxins, such as the human *Stx7* and the yeast (*S. cerevisiae*) VAM3 and PEP12 proteins (Wong *et al*., 1998; Teng *et al*., 2001).

### Syntaxin-deficient *Stentor* cells localize aberrant vacuoles at the cell surface

Having identified *Stentor* Stx as vital for post-wound cell survival, we next investigated protein function by perturbing the gene in native, unwounded cells. Syntaxin-deficient cells were characterized by the presence of aberrantly localized protrusions on the cell cortex, which we do not see in control cells (Fig. 3A). These aberrant structures were seen in cells targeted by RNAi constructs that targeted non-overlapping parts of the gene, Stx-C1 (residues 39-424) and Stx-C2 (residues 425-794) (Fig. 3A, Supplementary Table T1). Whereas control cells had a characteristic conical shape, syntaxin-deficient cells were often smaller and rounded.

**Figure 3.**
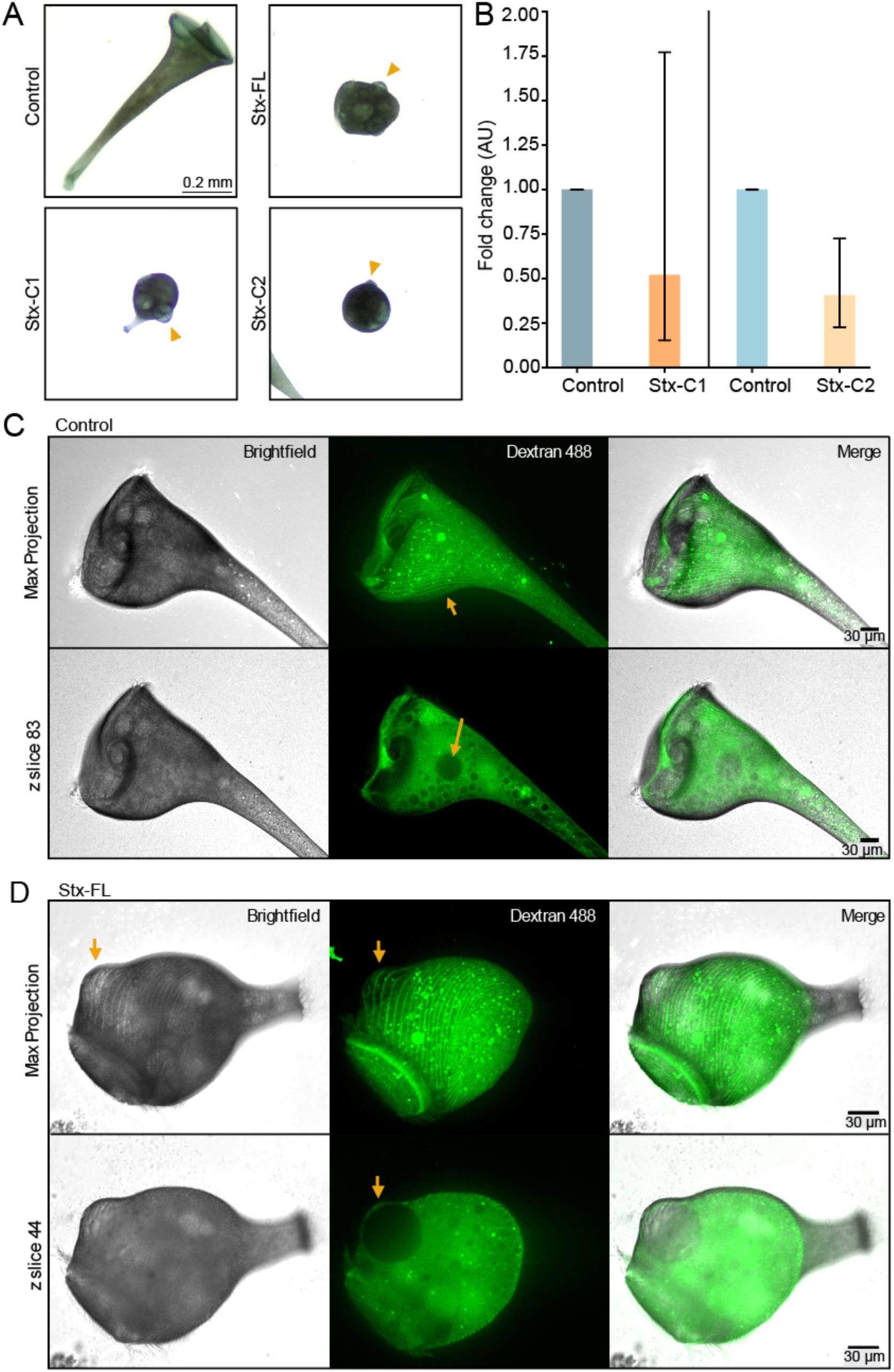
Syntaxin deficient cells aberrantly localize vacuoles. **(A)** Brightfield images of control (top left), Stx-FL(top right), Stx-C1 and C2 (bottom panels) depleted cells show aberrantly localized protrusions on the cell surface (yellow triangles) as a consistent phenotype between knockdown cells. **(B)** Fold change of gene expression in non-overlapping knockdown constructs is depicted with 95% confidence interval (error bars) of n=3 technical replicates. **(C)** Representative images of syntaxin knockdown cells injected with Alexa-488 dextran reveals that the vacuoles are found relatively interior (z slice 83 - bottom panels) of control cells (arrows) and cortical rows are not perturbed by their presence (top panel). **(D)** In Stx-FL knockdown cells, vacuoles (arrows) are found proximal to the cell surface with cortical microtubules curving around them.

We used qPCR to assess the extent of gene knockdown in cells targeted by the Stx-C1 and Stx-C2 constructs (see Methods). GAPDH transcripts were utilized for normalization (Slabodnick *et al*., 2014) (Fig. 3B). qPCR analysis revealed that syntaxin mRNA levels in cells transfected with the knockdown constructs (C1 and C2) were approximately 50% lower than in control cells (0.520x in cells targeted by the Stx-C1 construct and 0.404x for Stx-C2 - Fig. 3B, fold change and 95% confidence interval reported).

In order to determine whether the protrusions on the cell cortex in syntaxin-deficient cells were blebs of the plasma membrane pulling away from the cytoskeleton or vacuoles deforming the surface, we microinjected Alexa-488 labeled dextran (m.w. 10,000 Da) in control and knocked down cells. The dextran appeared to label the cortical microtubules (known as KM fibers; (Huang and Pitelka, 1973)) in some cells by an unknown mechanism. In control cells, vacuoles could be visualized as regions in the cytoplasm with low dextran fluorescence (Fig. 3C). These vacuoles were found towards the cell interior relative to the cortical microtubules, and the microtubules were unperturbed by the vacuoles. In contrast, in Stx-FL knockdown cells, we observed large vacuoles that were pushed against the cell surface, with the KM fibers curving around them (Fig. 3D). We confirmed this outwards bulging of the KM fibers using immunofluorescence of control and knockdown cells (Supplementary Fig. 3). Our results indicate the presence of membrane-bound vacuoles close to the cell-surface that are impermeable to dextran, and excludes the possibility that the structures are contiguous with the rest of the cytoplasm. The presence of unfused vacuoles at the cell-surface in syntaxin-deficient cells is consistent with the protein’s functions (Bennett *et al*., 1992; Rothman, 1994; Teng *et al*., 2001).

### Investigating the functional role of *Stentor* Stx

Our results indicated *Stentor* Stx was likely involved in vacuolar fusion at the plasma membrane. We hypothesized that deficiencies in vacuolar dynamics in Stx-FL RNAi cells would result in difficulties adapting to hyposmotic stress. In ciliates, hyposmotic stress increases the activity of the contractile vacuole, a specialized organelle involved in the excretion of water and ions by direct exocytosis at the plasma membrane (Cosgrove and Kessel, 1958; Allen, 2000; More *et al*., 2024). Contractile vacuole activity also involves internal vesicular dynamics that sustain the radial collection channels that feed into the central vacuole (Allen, 2000; More *et al*., 2024).

We placed unwounded control and Stx-FL RNAi cells in hypotonic media. While control cells were able to tolerate this osmotic challenge over 90 minutes, we observed accumulation of fluid into, and apparent lysis of large vacuoles in Stx-FL RNAi cells. We also observed the accumulation of secondary vacuoles in Stx-FL RNAi cells (Fig. 4).

**Figure 4.**
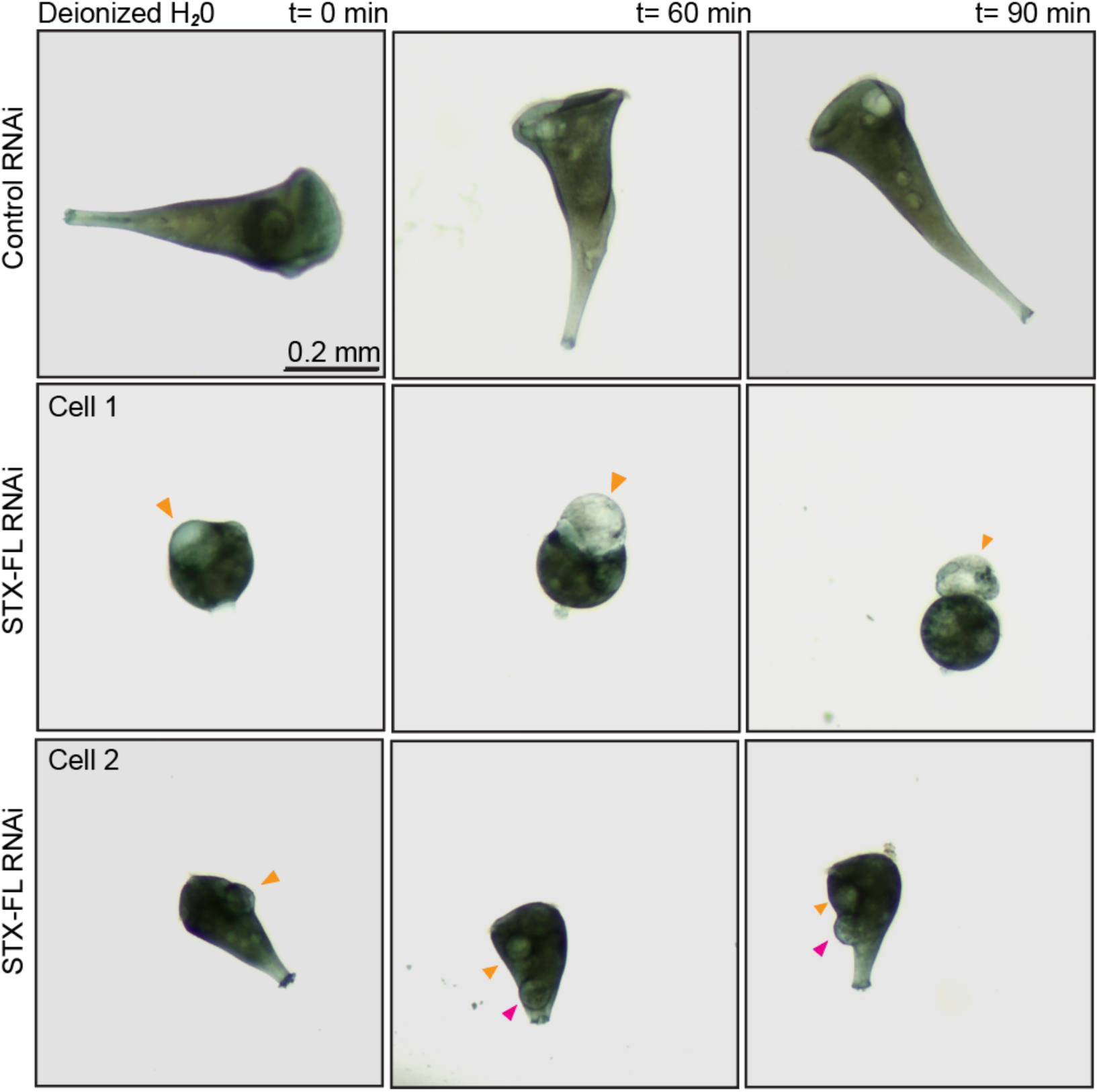
Syntaxin deficient cells are not as osmotically adaptable as control cells. Control (top) cells are able to endure the hyposmotic challenge of being placed in deionized water, whereas vacuoles in syntaxin-deficient cells (yellow arrowheads) undergo swelling (Cell 1 - middle panel) and apparent lysis (Cell 1 - right panel). Cells also may accumulate additional vacuoles (pink arrowheads, Cell 2, bottom panels). Representative images of n=3 independent experiments with 30 cells total.

The observed phenotype of Stx-FL RNAi cells suggests deficiencies in exocytic and vacuolar homeostasis mechanisms and offered us a plausible mechanism for their reduced viability post-wounding compared with control cells. When the cell membrane is ruptured during wounding, water can flow freely into the highly concentrated cytoplasm. After the cell has healed the wound and restored membrane integrity, it would be necessary to remove this water. One explanation for the Stx-FL RNAi phenotype would be an inability to excrete this excess water due to defects in contractile vacuole function.

Alternatively, syntaxins could potentially be involved in membrane patching immediately following wounding. A time course of cell survival post wounding revealed that all cells remained alive for at least 45 minutes post-bisection (Supplementary Fig. S4A,B). We observed the first events of cell death at 90 minutes. Qualitatively, cells became more vacuolated over time (Supplementary Fig. S4A), dying gradually with ∼75% cells surviving at 6 hours (consistent with our 12-hour survival statistic in Fig. 1). The fact that cells remain viable for so long post-wounding argues against a fatal defect in membrane patching.

### Wounding in an osmotically balanced medium provides Stx-FL with a survival boost

The inability of Stx-FL RNAi cells to recover from wounding-associated osmotic changes could explain their low viability. If so, then it should be possible to reduce the lethal effects of the knockdown by reducing osmotic water entry into wounded cells. Osmotic stabilizers such as sorbitol are utilized in the isolation of delicate structures such as protoplasts to prevent water influx (Yang *et al*., 2024). We asked if wounding the cells in a solution containing such osmotic stabilizers would provide syntaxin-deficient cells a survival boost.

Intracellular osmolarity has been found to be within the ranges of 75-300 mOsm (Milo *et al*., 2010), BNIDs 112732 & 104055. Qualitative experiments indicated that unwounded control and Stx-FL RNAi tolerated isosmotic conditions, accompanied by cellular and vacuolar shrinkage of knockdown cells (Supplementary Fig. S4C). We empirically determined a concentration of sorbitol (75 mM) which would preserve cell viability overnight. After acclimatizing the cells in this medium for 30 minutes, we conducted a wounding survival assay in 75 mM sorbitol.

To test our hypothesis, we modified our experimental setup used to wound the cells. We observed that sorbitol-acclimated Stx-FL RNAi cells were injured but not bisected by the guillotine employed in our original survival assay comprising 90° blades (Fig. 1). We speculated that this phenomenon was caused by cell-shrinkage in sorbitol solution.

To address this issue, we transitioned to a scaled-down guillotine where the channel was reduced by a factor of 2^1/3^ in each linear dimension to better match the reduced cell volume of Stx-FL sorbitol treated cells which was approximately half of the control cells (see Methods; data available upon request). This scaled-down device incorporated angled blunt-to-sharp (bts) blades that have been empirically shown to bisect smaller cells efficiently (Koppaka *et al*., 2021). We also used the 2^1/3^ bts guillotine to bisect Stx-FL RNAi cells in PSW. Stx-FL RNAi PSW and sorbitol cells had comparable cell volumes and we wanted to maintain consistent bisection conditions between the Stx RNAi cells. Normal-sized control cells (in PSW or sorbitol) were bisected using the standard-sized bts guillotine and showed 100% survival in all experiments. Importantly, the flow velocity at the blade was similar in all devices to ensure a consistent severity of wounding (see Methods).

The assay was calibrated for the new blades and devices by measuring cell survival in normal cell media (PSW). Stx-FL cells once again fared worse than control cells, with the fraction of surviving cells being 0.448 for knockdown versus 1.0 for control cells. This was consistent with our observations in Fig. 1 and our empirical evidence that bts blade provided a “gentler” bisection (Koppaka *et al*., 2021). Acclimatization in 75 mM sorbitol provided a moderate survival boost for Stx-FL RNAi cells (fraction surviving increasing from 0.448 to 0.761 in PSW versus sorbitol), thus partially rescuing the post-wound survival of RNAi cells (Fig. 5B). A Fisher’s exact test revealed that the difference in survival was statistically significant (p = 0.001) (Fig. 5C).

**Figure 5.**
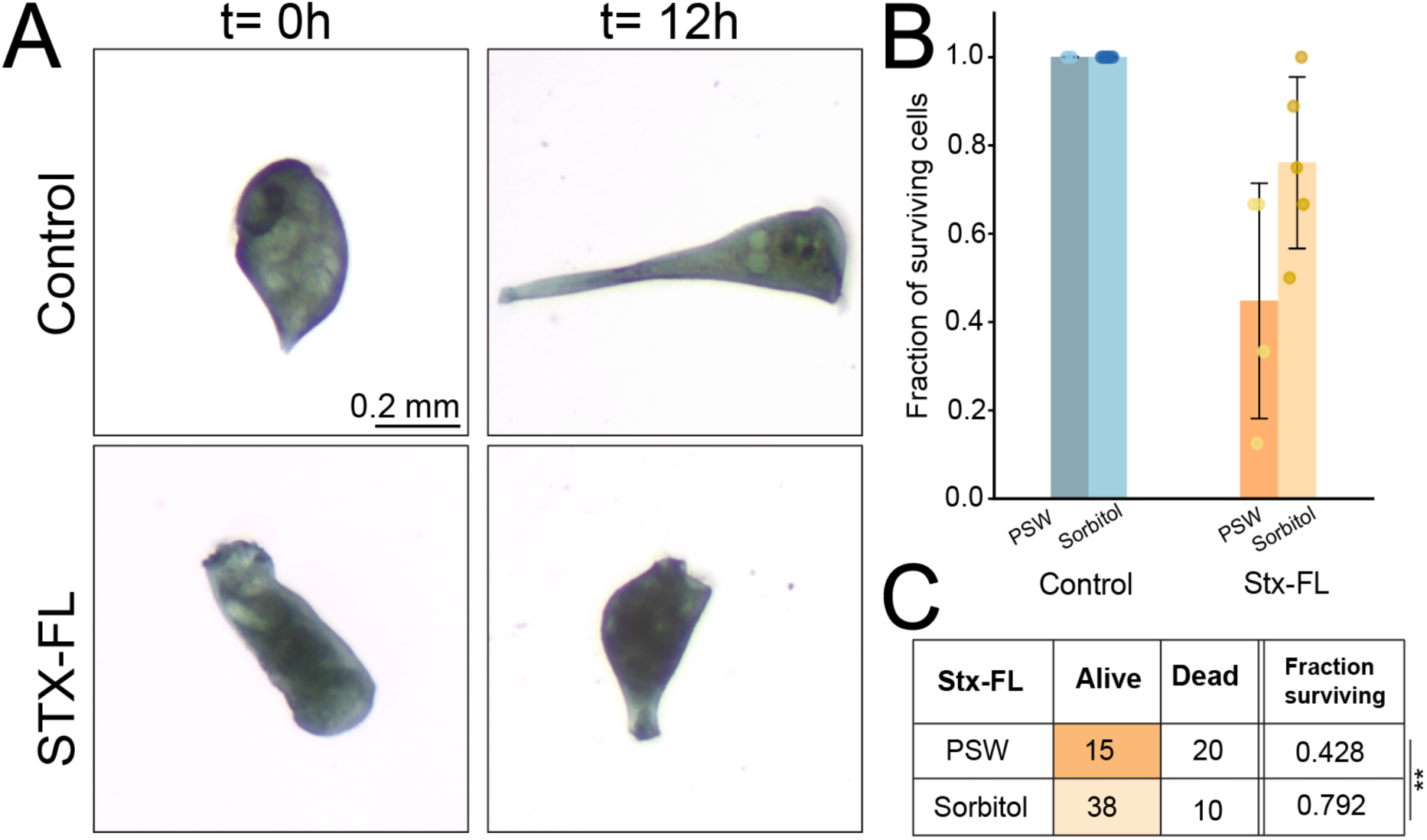
Bisection in osmotically-balanced conditions provides a mild survival advantage over wounding in spring water. **(A)** Control (upper) and Stx-FL RNAi (lower panels) were wounded in 75 mM sorbitol. Cells were visualized immediately after bisection (left panels) as well as 12h post wounding (right panels). **(B)** The fraction of surviving cells was quantified. Individual biological replicates are denoted as filled circles. **(C)** The outcome of cell survival compared between Stx-FL RNAi cells in control (PSW-dark yellow) or isosmotic (sorbitol -light yellow) media indicated by the total number of cell fragments across n=4 biological replicates in each condition is statistically significant by a two-sided Fisher’s exact test (p = 0.001).

In conclusion, we have identified a novel *Stentor*-like syntaxin protein that is essential for cell-survival post wounding. This discovery emerged from a targeted genetic screen that employed a microfluidic guillotine to enable high-throughput bisection of cells in a fast and reproducible manner (Blauch *et al*., 2017; Slabodnick, 2019). High-throughput microfluidic wounding devices represent engineering advances over traditional microsurgery methods that when combined with automated image analysis and the use of tracer dyes or small molecules, can be utilized for versatile applications such as extracting morphometric information or conducting small-molecule screens.

Cells deficient for the syntaxin gene died at significantly higher rates than control cells post-bisection. We discovered that the syntaxin protein contains canonical features of other homologs including syntaxin and SNARE-domains and a conserved glutamine residue. A putative palmitoylation site at the C-terminal domain may indicate a mechanism of membrane localization for the protein. The importance of these residues in plasma membrane-localized syntaxins represents an interesting future avenue of research.

Knockdown of Stecoe_24534 in unwounded cells led to aberrant vacuole localization and deficiencies in osmotic regulation of cells. We were able to partially rescue the post-wounding survival defect by wounding cells in osmotically balanced conditions. Our observations are consistent with a model in which water can flow into the cytoplasm rapidly when the membrane is broken, and must then be removed once membrane integrity is restored. In ciliates, as with other fresh-water protists, a contractile vacuole is required to expel water from the cytoplasm (More *et al*., 2024). In *P. tetraurelia*, syntaxin-2 (Syx2) has been found to localize to the contractile vacuole, and other syntaxins such as syntaxin-1 (Syx1) isoforms localize to the plasma membrane (Kissmehl *et al*., 2007). *Stentor* syntaxin could either be directly involved in contractile vacuole function like Syx2 or play an upstream role in the osmoregulation pathway by mediating fusion of internal vacuoles that eventually merge with the contractile vacuole. Further details of the molecular process remain to be elucidated and are outside of the scope of this current work.

Future work might include detailed characterization of the role of syntaxin in osmotic regulation at the contractile vacuole and associated structures. Syntaxin-like proteins have been found to be important during osmotic stress in soybean however they seem to act upstream of the abscisic acid signaling pathway and the details at the cell-biological level are unclear (Chen *et al*., 2019).

In the current study, candidate genes were knocked down using traditional RNAi by feeding. However, RNAi perturbation by microinjection has recently been shown to be a more effective method for genetic perturbation in *Stentor* (Kuecks *et al*., 2025) and represents a future novel methodology for the study of syntaxin function in *Stentor*.

Like single-celled freshwater ciliates such as *Stentor*, osmotic effects are also important during wound healing in multicellular organisms. For example, it has been shown that cell swelling due to influx of water and drop in interstitial osmolarity stimulates wound healing in zebrafish (Enyedi *et al*., 2013; Gault *et al*., 2014). Hypotonic stress causes phospholipase 2 to translocate to the nucleus which in turn mediates the release of the inflammatory mediator arachidonic acid and recruits immune cells. In humans, extracellular osmolarity can vary during normal physiological processes as well as in pathological conditions such as hyperosmolar hyperglycemia and dilutional hyponatremia (Olmstead, 1966; Pasquel and Umpierrez, 2014). Our results suggest that such conditions might place additional stress on wound-repair pathways (potentially including syntaxins or other trafficking regulators) and compromise the ability of mechanically active cells such as skeletal or cardiac muscle cells to survive wounding. Although there exist fundamental differences in osmotic regulatory mechanisms in animals and *Stentor*, water transport may be a conserved aspect of wound recovery at the cellular level.

## Materials and Methods

### Stentor cell culture

*Stentor coeruleus* were cultured in Pasteurized Spring Water (PSW) (132458, Carolina Biological Supplies) as previously described (Lin *et al*., 2018; Zhang *et al*., 2021). Cells were fed *Chlamydomonas reinhardtii* grown in a liquid suspension culture. A new feeding culture was inoculated every 7-10 days as described (Lin *et al*., 2018).

### Cloning

The plasmid containing *Stentor* Stx (designated Stx-FL in the text) was generated by PCR (Radiant Inc.) using a sequence corresponding to nucleotide residues 34-794 of the coding sequences of the gene SteCoe_24534, a putative syntaxin protein. Non-overlapping RNAi constructs Stx-C1 and Stx-C2 corresponded to residues 39-424 and 425-794, respectively. Information about the gene sequences in the study is contained in Supplementary Table T1. Primer sequences are described in Supplementary Table T2. Genomic DNA was isolated from *Stentor coeruleus* by using either a phenol chloroform extraction protocol or the Quick-DNA/RNA Microprep Plus kit (Zymo Research - manufacturer’s instructions). Briefly, for the phenol chloroform protocol, ∼200 cells were lysed into a buffer containing 100 mM Tris pH 9.0, 100 mM EDTA and 1% SDS. After a 30 min incubation at 70°C, potassium acetate was added to a final concentration of 0.7 M. To the supernatant, one volume of phenol/chloroform/isoamyl solution was added (Sigma Aldrich). DNA was precipitated using 0.7 volumes of isopropyl alcohol, washed with 70% ethanol and resuspended in autoclaved water after drying. Genes were cloned from genomic DNA using Phusion Polymerase (New England Biosciences, manufacturer’s instructions) and subcloned into the previously utilized pPR-T4P vector system using ligation independent cloning (Aslanidis and de Jong, 1990; Adler and Alvarado, 2018; Slabodnick, 2019). Plasmid products were sequenced by Elim Biopharma.

### RNA interference by feeding

RNA interference (RNAi) by feeding was carried out as described (Slabodnick *et al*., 2014; Slabodnick, 2019). Briefly, HT115 *E.coli* were transformed with plasmids containing genes of interest, grown and induced with 1 mM IPTG. Bacteria were pelleted after 3 ½ hours, washed with PSW and flash frozen in aliquots. An aliquot of frozen food equivalent to 200 µL of bacterial culture was fed per day to 100 starved *Stentor* cells (per condition). If using fewer cells, the food was scaled down proportionally. Phenotypes described were observed starting on day 6-7 and experiments were carried out on days 7-10. Cells were selected visually, using a dissection microscope, for experiments based on their small size, rounded morphology and presence of unfused vesicles at the cell-surface. A total of ∼15-30% percent of cells yielded a visual phenotype corresponding to roughly 30-60 knocked down cells per 200 *Stentor*.

### Microfluidic devices

The 90 ° radial guillotine was fabricated in poly(dimethylsiloxane) (PDMS) using standard photolithography as extensively described previously (Blauch *et al*., 2017; Zhang *et al*., 2021). The blunt-to-sharp guillotines were fabricated as described previously (Koppaka *et al*., 2021). Briefly, channel mold designs were first created in SolidWorks (Dassault Systèmes), then fabricated 2-photon polymerization in IP-S photoresist on a silicon wafer. This process was performed using a 25X immersion objective (NA 1.4) in the Nanoscribe Photonic GT system (Nanoscribe, GmbH). Because of poor adhesion of the IP-S photoresist to the silicon substrate, only a single PDMS device was cast from the original mold. The cast PDMS device was then used to make a polyurethane replica. The polyurethane replica was used as the mold for subsequent devices. A smaller device scaled down by 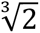 (with twice the number of parallel guillotines) was used for smaller syntaxin-deficient cells (only for experiments in Fig. 5).

### Osmotic experiments

For the hypotonic experiments in Fig. 4, cells were washed into and imaged in deionized water using the Zeiss Stemi 508 dissection microscope with the Axiocam 208 color camera for 0-90 min. For experiments in Fig. S4, a 100 mM and 75 mM concentration of sorbitol was utilized (dissolved in PSW) where indicated.

### Survival Assay

Cells were collected and washed at least 2x with PSW to reduce stickiness caused by RNAi food. For experiments in Fig. 1 and Fig. 1S, cells were injected into the radial microfluidic guillotine device using a fluidic pump at 1.4 cm/s per blade using a 10 cm-long outlet tubing and a fixed volume elution of 0.5 ml, as described previously (Zhang *et al*., 2021). Each repeat survival assay provided us with a percent cell survival, so though we tested roughly 40-50 cell fragments on average, the total n ∼ 4 (indicated in each figure caption). Due to the fact that our control cell survival was often 100% for every repeat, we couldn’t verify normality of our data and therefore used the non-parametric Mann-Whitney test to measure statistical significance in Fig.1.

For the experiments shown in Fig. 5, two media conditions were used, and microfluidic guillotines with angled blunt-to-sharp (bts) blades were used.

For the sorbitol experiments, control and Stx-FL RNAi cells were first acclimatized in 75 mM sorbitol in PSW for 30 minutes.

- Control cells were cut using a normal-sized 4-blade guillotine at 1.4 cm/s per blade.
- Stx-FL RNAi cells were cut using a guillotine that was scaled down in each linear dimension of the channel (height and width) by a factor of 2^1/3^, but that had twice the number of blades (8) with an average velocity of 1.1 cm/s per blade.

Flow velocity at the blade relative to the cell diameter was similar between the two devices, ensuring a consistent severity of wounding. Once eluted from the microfluidic guillotines, viable cell fragments were immediately counted at t = 0 h using a dissection microscope (Zeiss Stemi 508) equipped with an Axiocam 208 color camera. Cells were counted again at t = 12 h, with cellular or ciliary movement serving as indicators of viability.

### Microinjection

Microinjection was carried out as previously described (Diaz *et al*., 2021; Diaz, 2024; Kuecks *et al*., 2025). Needles were pulled using a P-97 Flaming/Brown Micropipette Puller (Sutter Instruments). Needles were cut using a razor blade at an empirically determined distance to enable injection. 10 µl of injectant, 0.5 µg/µl or ∼ 50 µM fluorescently labeled Dextran Alexafluor 488 (Thermofisher, Catalog number D22910) in PSW was spun down in a 1.5 ml Eppendorf tube for approximately 1 minute, using a mini centrifuge, to remove clumps that may clog the needle tip. Needles were loaded with 0.5-1 µL of injectant approximately 5 minutes before injection. Approximately 50 (per condition) healthy control and knocked down *Stentor* cells were starved for 12 hours and placed in 1.5% methylcellulose solution to allow cells to shed their pellicle and enable easier injection. Microinjection was carried out using a Drummond Nanoject II system (discontinued) and monitored using a Zeiss Stemi 508 stereo microscope with a Transillumination 300 and dark field system.

### Live cell imaging

For images in Fig. 1C, 3A and 3B, cells were imaged in PSW using a Zeiss stereo microscope (Zeiss Stemi 508 with an Axiocam 208 color camera) in a 12 well-plastic or 24-well glass-bottomed plate. Microinjected cells were imaged using an inverted Nikon Ti2-E microscope equipped with a CREST X-Light V2 large field of view spinning disk, a Prime 95 25MM sCMOS camera, and a Celesta Light system. Fig. 3C and 3D were imaged on this system using a Nikon CFI Plan Apochromat Lambda S 25XC Sil objective. Live cells were mounted in methylcellulose 1.5% solution and imaged in a 9mm diameter, 120 µm thick spacer (grace bio labs SecureSeal) between a standard glass slide and number 1.5 coverslip. Images were segmented, cropped and registered using the OttoReg segmentation pipeline developed by U. Diaz (Diaz, 2024). Ottoreg software was written using MATLAB (Mathworks Inc.).

### Bioinformatics

Domain analysis of the protein sequence Stecoe_24534 was performed using InterPro (Blum *et al*., 2025). Domains were depicted using the Illustrator for Biological sequences and adapted using Adobe Illustrator (Xie *et al*., 2022). Multiple sequence alignments were obtained using the protein sequences of the indicated genes and Clustal Omega or MUSCLE for alignments (Edgar, 2004; Sievers *et al*., 2011; Sievers and Higgins, 2018). Sequence alignment in Fig. S2 was generated using Jalview (Waterhouse *et al*., 2009). Multiple sequence alignment depicted in Fig. S3 was generated using ggmsa (Zhou *et al*., 2022). Biopython was utilized in the generation of the heatmap (Cock *et al*., 2009). The phylogenetic tree was generated using IQ-TREE3, using the best-fit model and a bootstrap value of 5000 (Kalyaanamoorthy *et al*., 2017; Hoang *et al*., 2018; Wong *et al*., 2025). Trees were visualized using FigTree (http://tree.bio.ed.ac.uk/software/figtree/) and iTOL (Letunic and Bork, 2007, 2024). Some fragments of code were generated with the assistance of ChatGPT, accompanied with extensive testing and debugging (https://chat.openai.com/). Fasta files, multiple sequence alignment files and code will be made available upon request.

### RNA extraction and quantitative PCR (qPCR)

RNA was extracted from ∼50 *Stentor* cells (per condition) using the Quickprep DNA-RNA mini kit (Zymo) using manufacturer’s instructions. On-column DNAse treatment was carried out at 1.5x the recommended volume (60 µL) and for 40 min instead of 15 min. RNA concentration was assayed using a ThermoScientific Nanodrop 2000.

Primers used for qPCR are described in Supplementary Table T2. RNA template was diluted to ensure roughly equal amounts per reaction. qPCR was carried out using a BioRad CFX-Connect thermocycler, using the iTaq™ Universal SYBR® Green One-Step Kit as previously described (Slabodnick *et al*., 2014) with an annealing temperature of 54°C. Controls lacking reverse transcriptase enzyme and template and a melt-curve were run at the end to ensure consistency between reactions and to verify the quality of RNA and product. GAPDH was used as a reference gene (Slabodnick et al, 2014). A standard curve for each primer pair was generated with an empirically determined threshold fluorescence level. Three replicates were run. The standard curve was used to determine the Cq and DNA amounts in ng for each condition. We obtained an average DNA amount in ng for control and knockdown conditions for both target (syntaxin) and reference (GAPDH) genes with associated standard error. Fold change was calculated by normalizing to the amount of reference gene in each condition as follows:

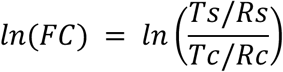

where T: target, R: reference, s: sample (knockdown), c: control.

Error was propagated using the formula:

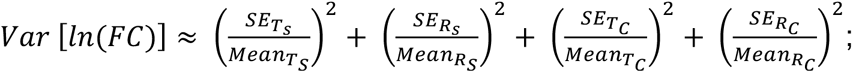 for independent samples. SE: Standard Error, Var: Variance. Confidence interval = 1.96 ± SE.

### Data analysis, images and figures

Data was graphed using Python packages numpy, matplotlib and seaborn (Hunter, 2007; Harris *et al*., 2020; Waskom, 2021). Other tools used were GraphPad online tools and Microsoft Excel. Statistical tests used are indicated in the figure legends. For Fig 1., control cell survival was often close to 100%, leading to a skewed distribution (by Shapiro-Wilk test for normality), so a non-parametric Mann-Whitney U-test was used. For Fig 5., due to the fact that we were measuring only a ∼1.5x increase of Stx-FL RNAi survival from PSW into sorbitol and we had a small experimental n (n=4), we pooled our cell survival data across biological replicates. This served to increase our statistical power as our experiments were unpaired, independent trials of cell survival with two possible outcomes (alive or dead). Fiji was used to generate images (Schindelin *et al*., 2012). Dynamic range of images (using min-max) was adjusted using Fiji for aesthetic quality. No quantification of data from modified images was carried out. Confocal images were scaled similarly to facilitate comparison. Non-similar modifications are indicated in the caption. Brightfield images were adjusted to enable visualization of cells against the background. Figures were made using Adobe Illustrator.

## Acknowledgements

Confocal imaging was performed at the Cell Sciences Imaging facility at the Stanford School of Engineering Shriram Center. qPCR was performed at the UCSF CAT, supported by UCSF PBBR, RRP IMIA, and NIH 1S10OD028511-01 grants. Thanks to Rebecca McGillivary, Connie Yan and Ashley Albright for invaluable assistance regarding *Stentor* protocols. Thanks to the Moumita Das lab (RIT) for insights and comments on the project. The Tang Lab (Stanford University) and Marshall Lab (UCSF) provided extensive feedback regarding the project. Thanks to Asst. Prof. Ranjani Murali (UNLV) comments regarding multiple sequence alignment and phylogeny. This work was supported by NSF Awards 2317442 and 1938109 to SKY and WFM, and by the Center for Cellular Construction, which is a Science and Technology Center funded by the National Science Foundation (NSF Award: DBI-1548297). This work was also supported by NIH grant R35 GM130327 to WFM. Device fabrication was performed in the Stanford Nano Shared Facilities (SNSF) and the Stanford Nanofabrication Facilities (SNF).

## Author contributions

The study was conceptualized by AVN, SKY and WFM. AVN, UD and KSZ carried out the experiments, collected and interpreted data. AVN and UD made the figures. AVN wrote the manuscript, with inputs from UD, SKY and WFM.

## Supplementary information

## Supplementary figures

**Figure S1.**
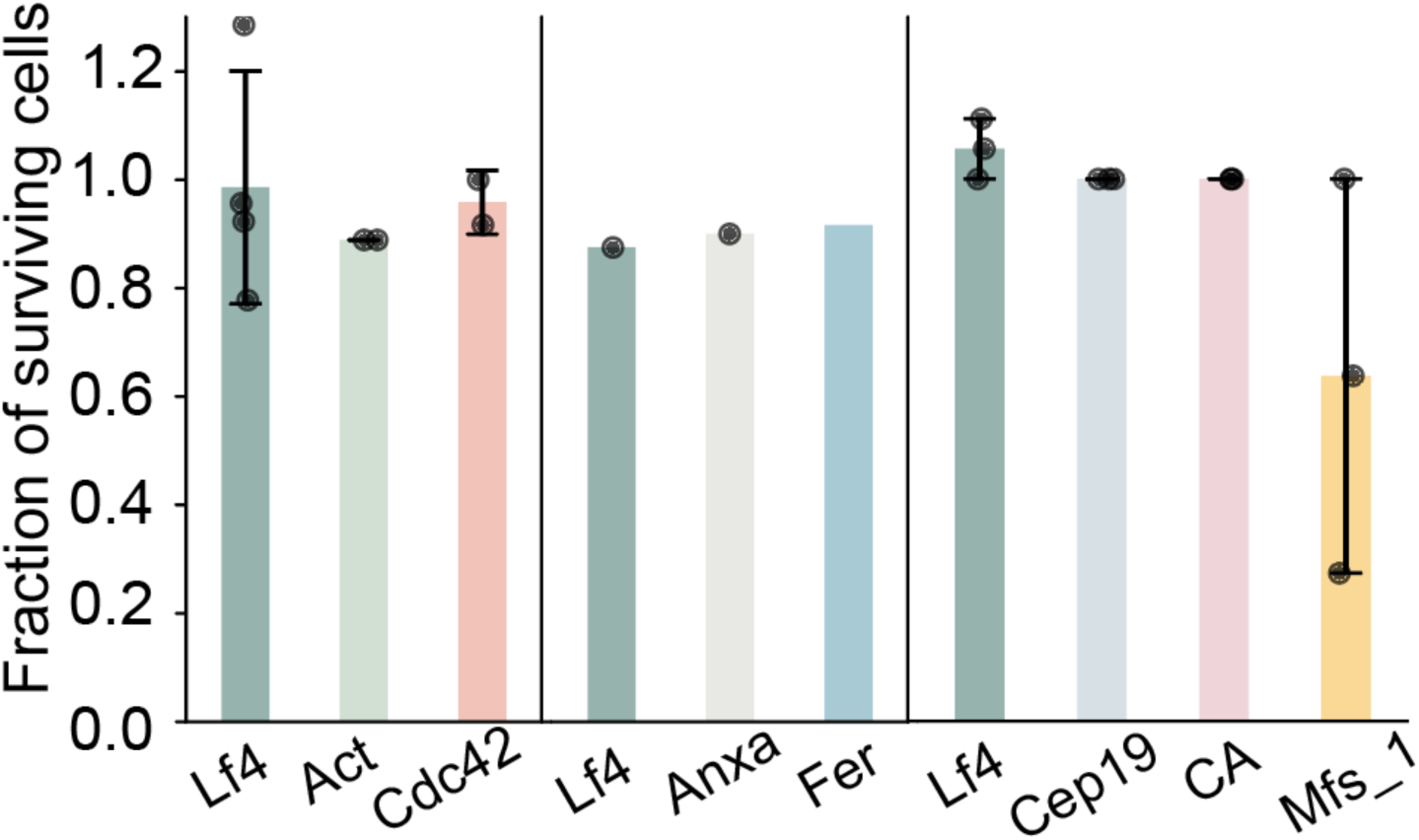
Survival assay of *Stentor* perturbed for candidate wound-healing genes. Genes indicated are as follows - Lf4 indicates a previously established control. Act - actin (Stecoe_7446), Cdc42 (Stecoe_21574), Anxa - annexin (Stecoe_18601). Fer - C2-domain containing protein-ferlin (Stecoe_31310), Cep19 - Stecoe_12911, CA-carbonic anhydrase protein (Stecoe_37123), Mfs_1-major facilitator superfamily 1 domain protein (Stecoe_168). Dots indicate individual experiments (biological replicates). Error bars represent S.D.

**Figure S2.**
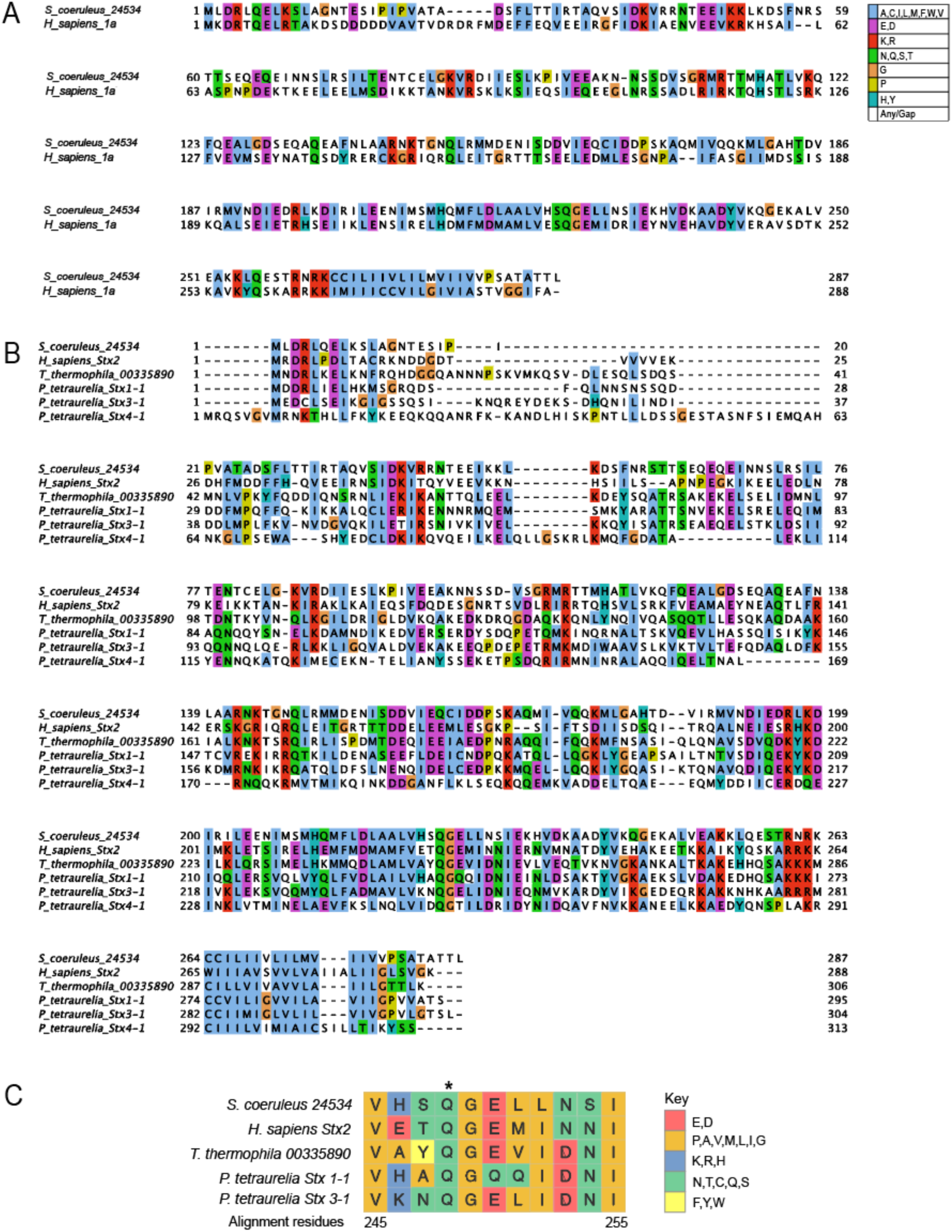
Multiple sequence alignment of syntaxin proteins. **(A,B)** Species and genes are indicated in the figure. Amino acids are colored using the Clustal scheme, key denoted. **(C)** Multiple sequence alignment of close homologs of *Stentor coeruleus* syntaxin protein 24534 in *Homo sapiens, Tetrahymena thermophila* and *Paramecium tetraurelia* identified by BLAST show the degree of amino acid conservation around the Q glutamine residue (*).

**Figure S3.**
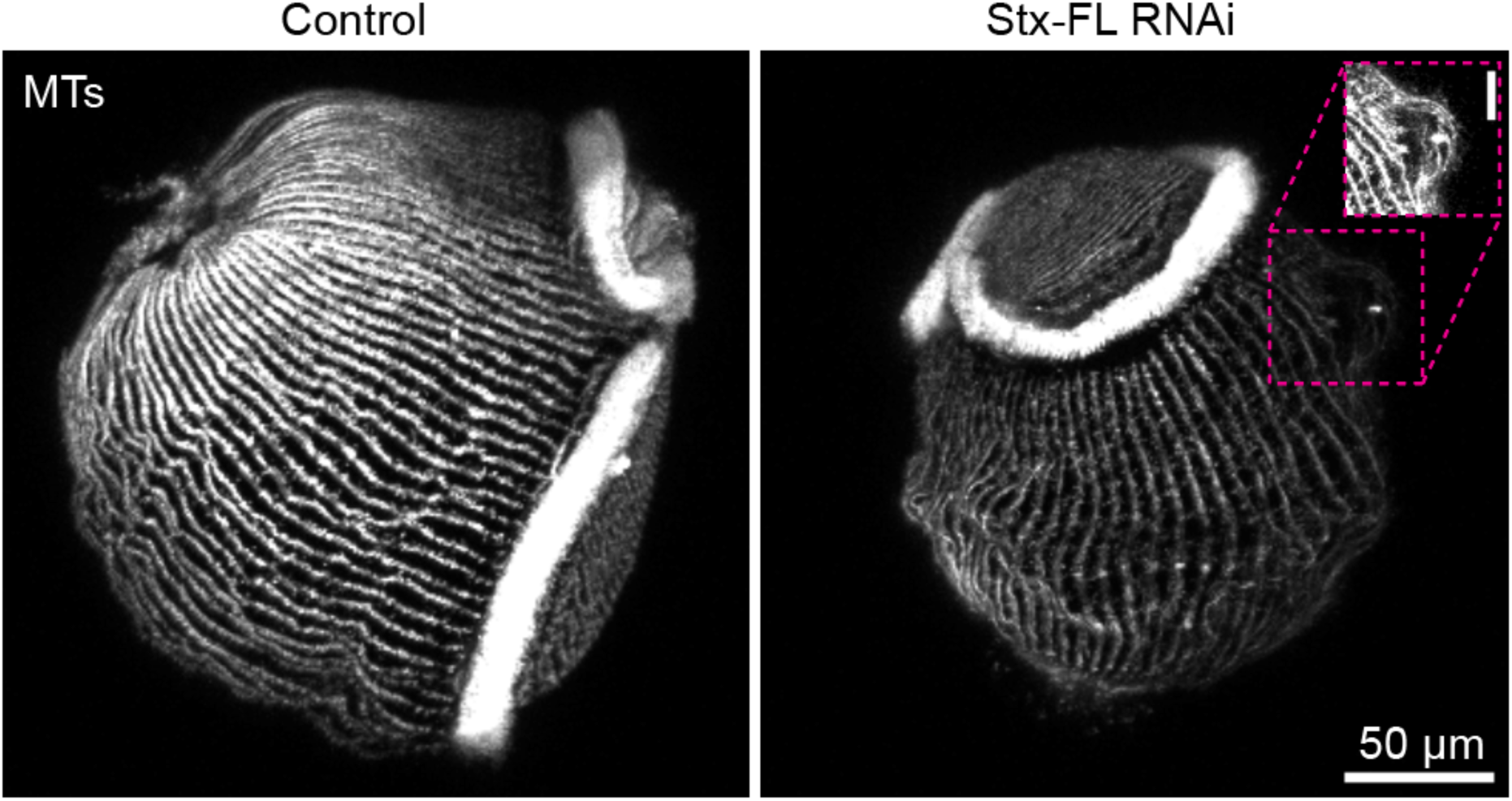
KM fibers curve around surface-level vacuoles in Stx-FL RNAi but not in control cells. Immunofluorescence of acetylated tubulin shows evenly spaced KM fibers with no aberrant curvature in control (left) panels and KM fibers curving around a vacuole (magenta box) in knockdown cells. Inset represents boxed area with enhanced dynamic range enabling visualization of the curved KM fibers. Scale bar in inset 10 µm. Representative images are from n=2 biological experiments.

**Figure S4.**
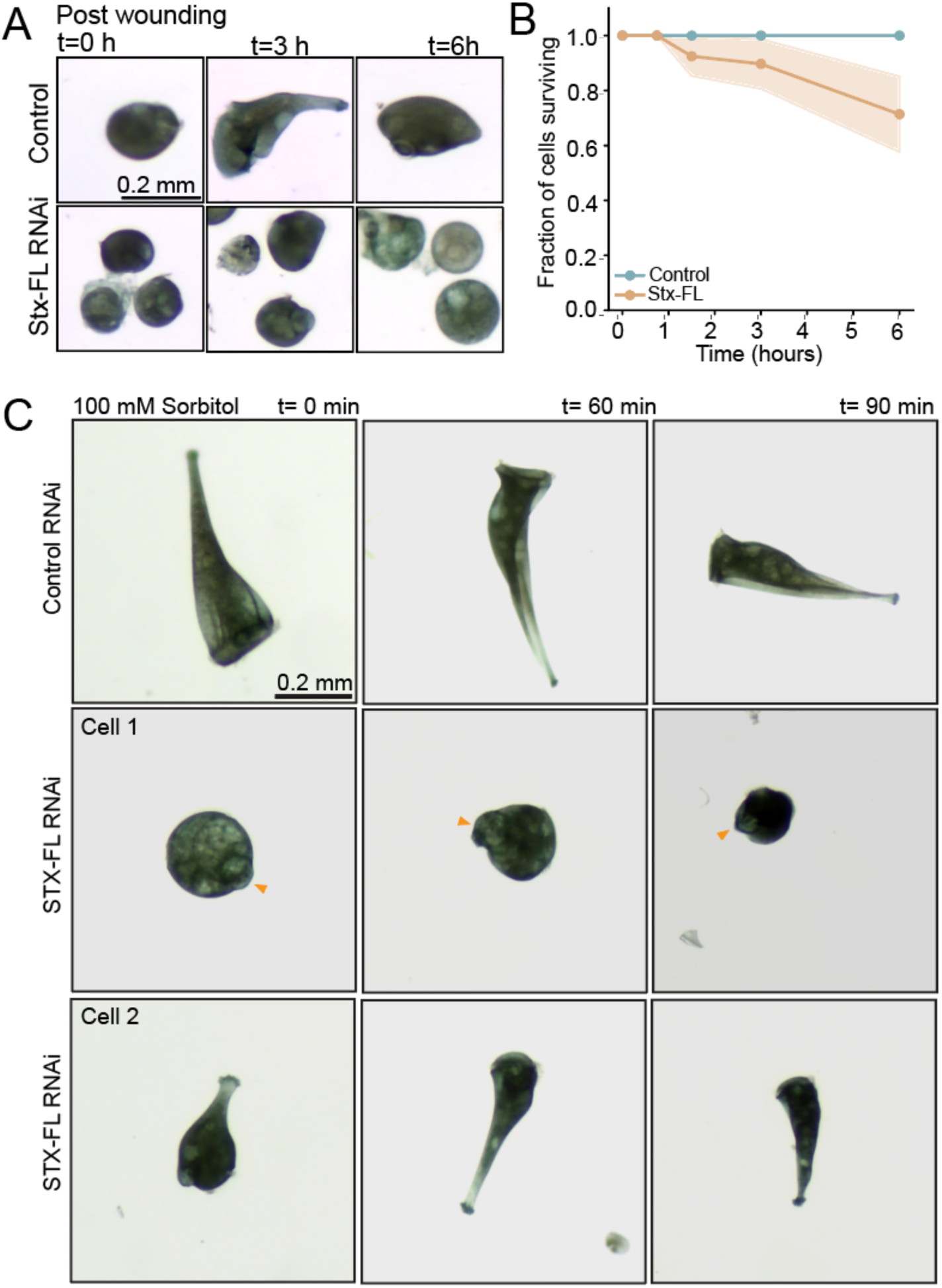
Mechanism of Stx-FL cellular deficiencies. **(A)** Brightfield images of control and Stx-FL RNAi cells at various time points post wounding (indicated) show that Stx-FL RNAi but not control cells look increasingly vacuolated over time. **(B)** Graph depicting the timescale of survival of control (blue dots and lines) and Stx-FL RNAi (yellow) cells. Line represents mean and shaded region represents SD of n=3 biological replicates, average 10 fragments per experiment. **(C)** Brightfield images depicting control and Stx-FL RNAi cellular response to osmotically-balanced conditions (100 mM sorbitol) reveals shrinkage of the vacuole in cell 1 (yellow arrowheads) and a return to relative conical shape in cell 2.

**Supplementary table T1.** Information about genes in the study.

| Gene/Protein | Functional significance/References | Stentor gene | Residues cloned (bases) | Primers Used (See T2) |
| --- | --- | --- | --- | --- |
| Stx_FL |  | Stecoe_24534 | 34-794 |  |
| Stx_C1 |  | Stecoe_24534 First half | 39-424 | Stx_c1_f/r |
| Stx_C2 |  | Stecoe_24534 Second half | 425-794 | stx_c2_f/r |
| Actin Domain Protein | Purse-string closure (Sonnemann and Bement 2011; Abreu-Blanco et al. 2011) | Stecoe_7446 | 12-1002 |  |
| Cdc42-like | Purse-string closure (Benink and Bement 2005) | Stecoe_21574 | 437-1100 |  |
| Annexin-like | Calcium-regulated protein, membrane transport, membrane repair in combination with dysferlin (McNeil et al. 2006; Gerke et al. 2024) | Stecoe_18601 | 26-239 | 18601_f/r |
| Ferlin (C2 domain protein) | Homologous to muscle dysferlin, acts in vesicle fusion, membrane repair in combination with annexins (Bansal et al. 2003; Lennon et al. 2003) | Stecoe_31310 | 1811-3408 | 31310_f/r |
| Cep19 domain protein |  | Stecoe_12911 | 4-265 | 12911_f/r |
| Carbonic anhydrase domain protein |  | Stecoe_37123 | 68-570 | 37123_f/r |
| Major Facilitator Protein_1 |  | Stecoe_168 | 160-1170 | 168_f/r |

**Supplementary table T2.**
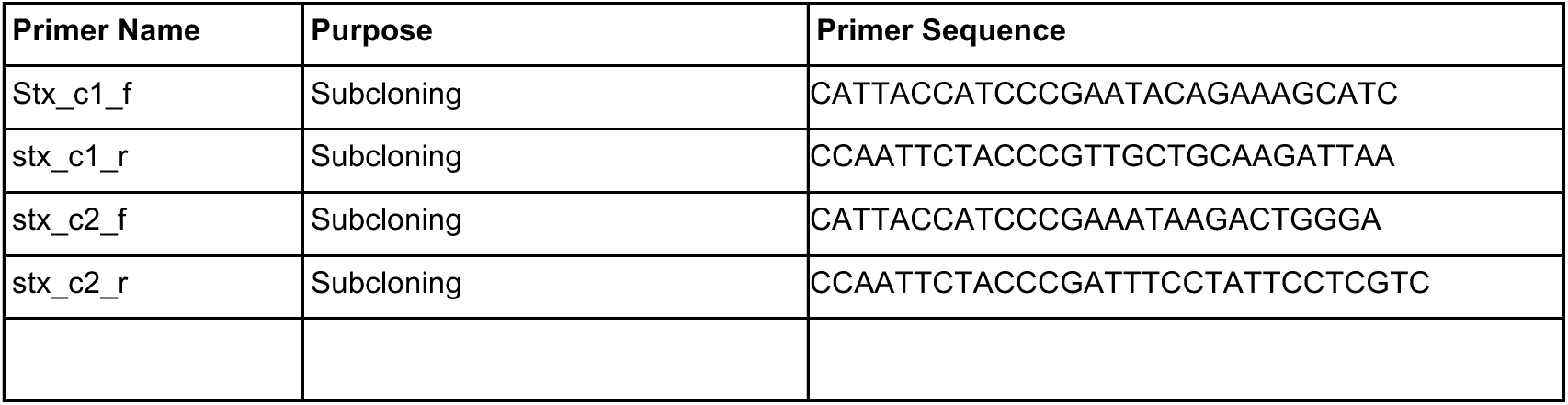
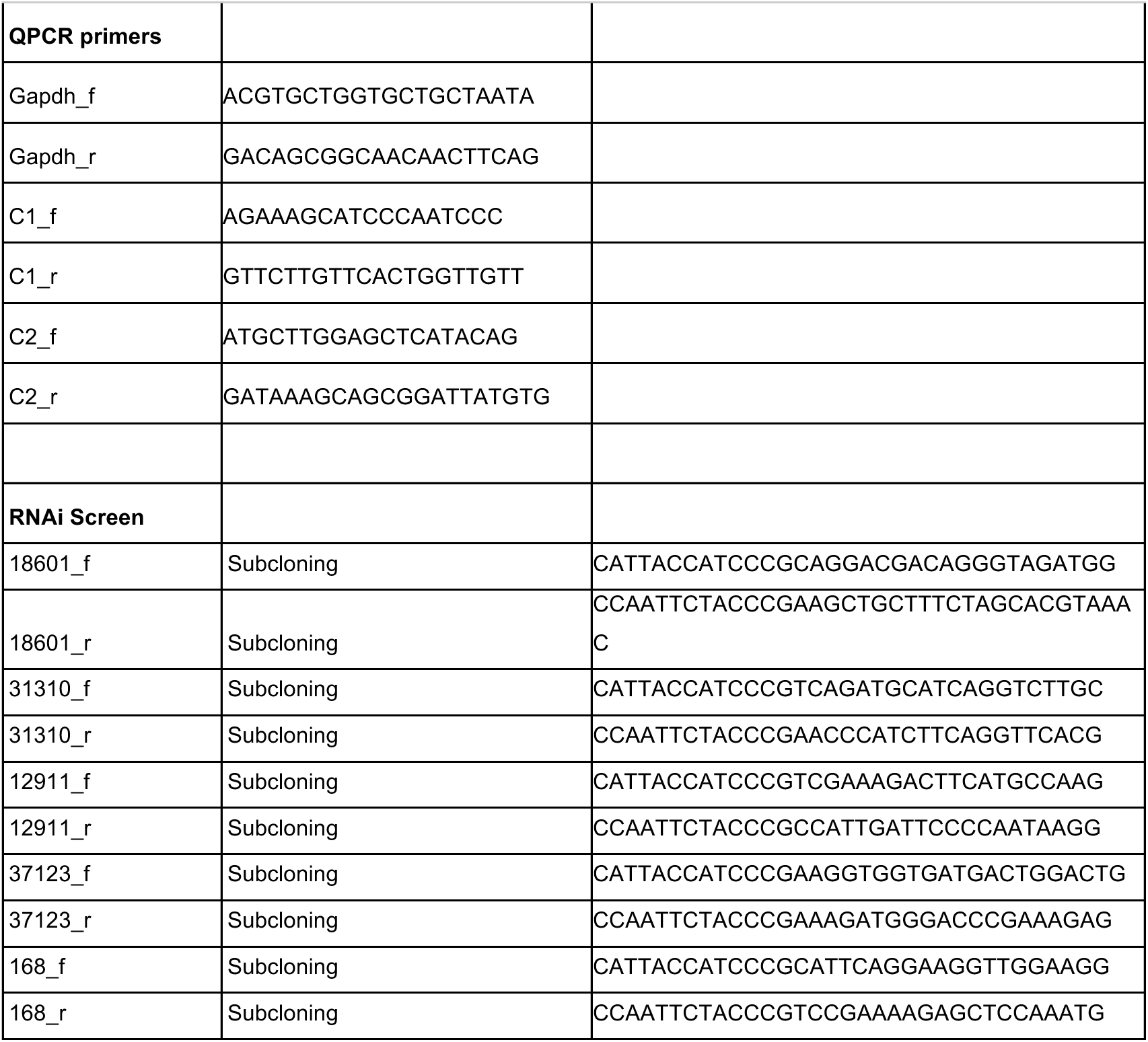
Primers utilized in the study.

